# Irradiated mesenchymal stromal cells induce genetic instability in human CD34+ cells

**DOI:** 10.1101/2020.10.30.361758

**Authors:** Vanessa Kohl, Oliver Drews, Victor Costina, Miriam Bierbaum, Ahmed Jawhar, Henning Roehl, Christel Weiss, Susanne Brendel, Helga Kleiner, Johanna Flach, Birgit Spiess, Wolfgang Seifarth, Daniel Nowak, Wolf-Karsten Hofmann, Alice Fabarius, Henning D. Popp

## Abstract

Radiation-induced bystander effects (RIBE) in human hematopoietic stem and progenitor cells may initiate myeloid neoplasms (MN). Here, the occurrence of RIBE caused by genotoxic signaling from irradiated human mesenchymal stromal cells (MSC) on human bone marrow CD34+ cells was investigated. For this purpose, healthy MSC were irradiated in order to generate conditioned medium containing potential genotoxic signaling factors. Afterwards, healthy CD34+ cells from the same donors were grown in conditioned medium and RIBE were analyzed. Increased DNA damage and chromosomal instability were detected in CD34+ cells grown in MSC conditioned medium when compared to CD34+ cells grown in control medium. Furthermore, reactive oxygen species and distinct proteome alterations, e.g., heat-shock protein GRP78, that might be secreted into the extracellular medium, were identified as potential RIBE mediators. In summary, our data provide evidence that irradiated MSC induce genetic instability in human CD34+ cells potentially resulting in the initiation of MN. Furthermore, the identification of key bystander signals, such as GRP78, may lay the framework for the development of next-generation anti-leukemic drugs.

## Introduction

Radiation therapy for neoplastic or non-neoplastic disorders may induce myeloid neoplasms (MN) in humans which are referred to as therapy-related myeloid neoplasms (t-MN) [1]. t-MN comprise the therapy-related cases of acute myeloid leukemia (t-AML), myelodysplastic syndromes (t-MDS) and myelodysplastic/myeloproliferative neoplasms (t-MDS/MPN) [1]. t-MN show genetic similarity to other high-risk MN [2, 3] and are associated with poor prognosis [4, 5]. Risk factors for the development of t-MN may include (a) inherited germline mutations in cancer susceptibility genes (e.g., *BRCA1*^wt/mut^), (b) acquired DNA damage in hematopoietic stem and progenitor cells (HSPC), (c) selection of pre-existing mutated hematopoietic clones (e.g., *TP53^wt/mut^*) and (d) alterations in the bone marrow stromal niche [6].

Irradiation may damage DNA directly in HSPC by interaction with DNA or indirectly by generation of free radicals [7]. In addition, irradiated mesenchymal stromal cells (MSC) may initiate radiation-induced bystander effects (RIBE) in HSPC potentially causing the development of t-MN [8, 9]. More precisely, RIBE describe ‘out of field’ effects of irradiation in non-irradiated cells which are mediated by genotoxic signaling factors released from nearby or distant irradiated cells [10]. RIBE may emerge as DNA damage (e.g., gene mutations, chromosomal aberrations, micronuclei, increased γH2AX foci), cell death (e.g., apoptosis, necrosis) and induction of cell survival mechanisms (e.g., adaptive response, DNA repair) [11–14]. Genotoxic signals, which mediate RIBE, are assumed to be initiated in irradiated cells by calcium fluxes [15] and mitochondrial metabolites [16–18]. Consecutively, signal transmission between irradiated and non-irradiated cells may occur by small molecules like nitric oxide (NO) [19], reactive oxygen species (ROS) [20], nuclear factor-kappa B (NF-kappa B) [18], and transforming growth factor beta-1 (TGFbeta-1) [21, 22], that pass through cell membranes and gap junctions [23, 24]. Finally, ROS generated by NADH oxidases [25] and unknown RIBE mediators may be induced in nearby or distant non-irradiated cells. RIBE have been studied so far in normal cells, cancer cells and in several *in vivo* model systems [9, 26–30]. While RIBE have been demonstrated in mouse HSPC [8, 9], RIBE have not been detected in primary human HSPC yet [31].

In summary, RIBE might be mediated in HSPC by genotoxic signaling from irradiated MSC and may account for a major pathomechanism in the initiation of certain MN. In contrast, the occurrence of RIBE in human HSPC has never been verified and genotoxic signaling factors are unknown yet. Therefore, our study was designed to analyze RIBE in CD34+ myeloid progenitor cells by immunofluorescence microscopy of γH2AX (as a readout of DNA damage), by analysis of G-banded chromosomes (for detection of chromosomal instability (CIN)) and by luminescence plate reading of cell viability. Furthermore, ROS and proteome alterations were assessed in irradiated MSC, MSC conditioned medium and CD34+ cells grown in MSC conditioned medium for the identification of potential genotoxic signaling factors.

## Materials and methods

### Femoral head preparation

This study was approved by the Ethics Committee II, Medical Faculty Mannheim, Heidelberg University. Procedures were performed in accordance with the local ethical standards and the principles of the 1964 Helsinki Declaration and its later amendments. Written informed consent was obtained from all study participants. Femoral heads of 12 patients with coxarthrosis (7 females, 5 males, mean age: 69 years) undergoing endoprothetic surgery were collected (Table 1). The bones were broken into fragments and incubated for 1 hour at 37 °C in phosphate-buffered saline (PBS) supplemented with 1 mg/ml collagenase type I (Thermo Fisher, Waltham, US). The supernatants were filtered through 100 μm pores of a cell strainer (Greiner Bio-One, Kremsmünster, Austria). MSC were grown from the fragments retained in the cell strainers in serum-free StemMACS MSC Expansion Media XF (Miltenyi Biotec, Bergisch Gladbach, Germany) supplemented with 1% penicillin/streptomycin. Adherent MSC were expanded in T175 flasks in a humidified 5% CO_2_ atmosphere at 37 °C and passaged at 80% confluency. Furthermore, CD34+ cells were enriched from the filtrates by Ficoll density gradient centrifugation and magnetic-activated cell sorting using CD34 antibody-conjugated microbeads (Miltenyi Biotec). CD34+ cells were grown in serum-free StemSpan SFEMII (Stemcell Technologies, Vancouver, Canada) supplemented with StemSpan Myeloid Expansion supplement (SCF, TPO, G-CSF, GM-CSF) (Stemcell Technologies) and 1% penicillin/streptomycin in a humidified 5% CO_2_ atmosphere at 37 °C.

**Table 1.**
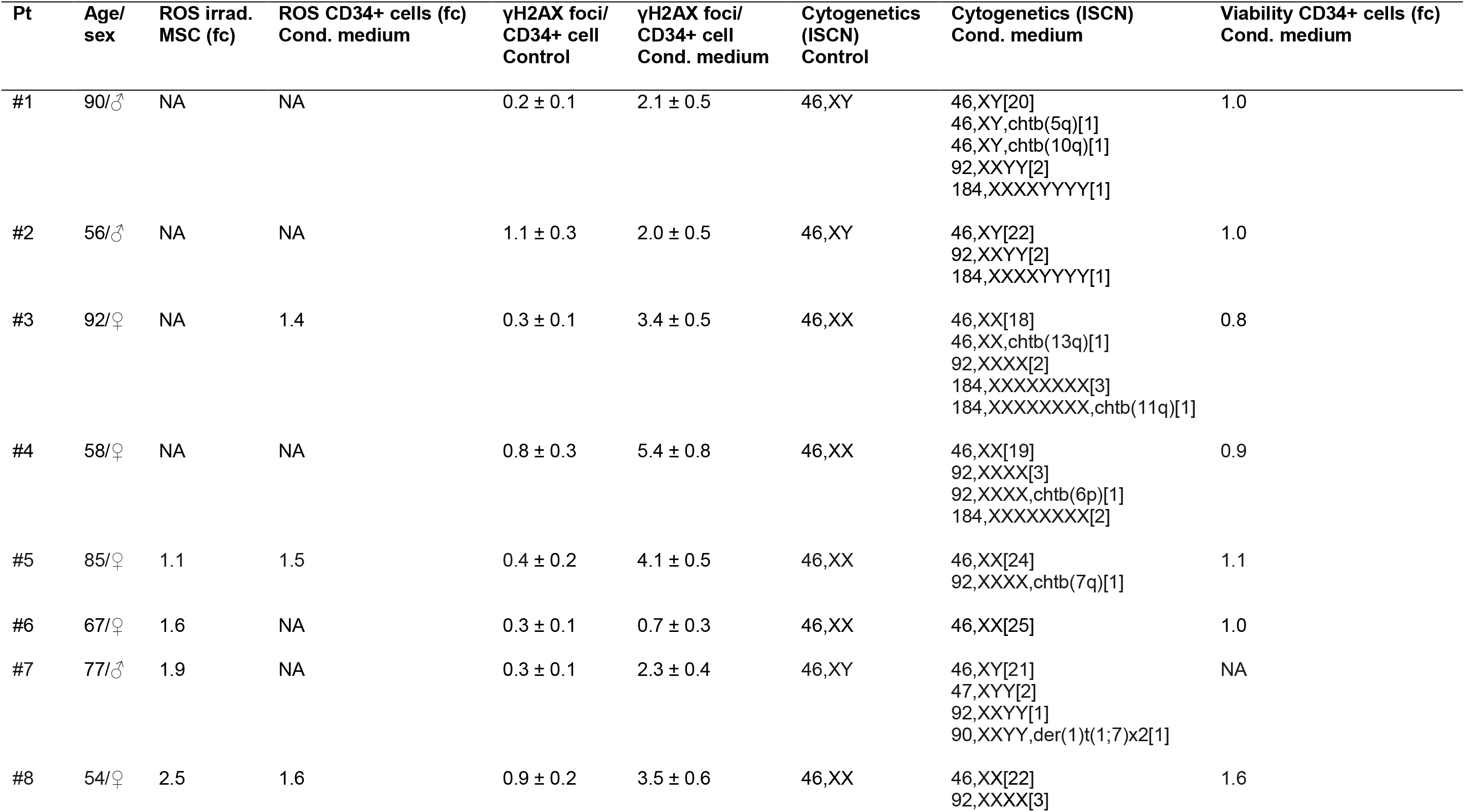
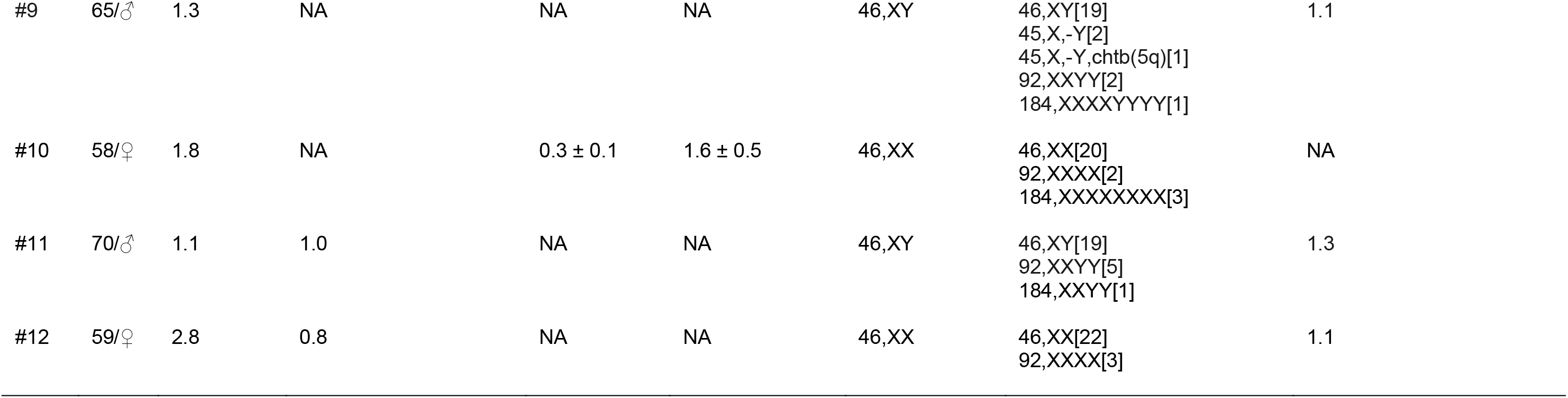
Radiation-induced bystander effects in CD34+ cells. *fc* fold change; *ISCN* international system for human cytogenetic nomenclature; *NA* not assessed; *Pt* patient; *ROS* reactive oxygen species.

### Preparation of MSC conditioned medium

MSC were grown in T175 flasks until reaching 80% confluency. MSC were rinsed in PBS before fresh serum-free StemSpan SFEMII was added. Afterwards, MSC were irradiated with 2 Gy of 6 MV x-rays in a Versa HD linear accelerator (Elekta, Stockholm, Sweden), while control MSC were not irradiated. Medium was conditioned by the irradiated and nonirradiated MSC for a period of 4 h incubation at 37 °C to generate MSC conditioned medium and control medium, respectively. The media were centrifuged at 1,200 rpm for 10 min, and supernatants were stored at −20 °C.

### RIBE analyses

RIBE were analyzed in CD34+ cells at day 6 after culture for 3 days in untreated medium followed by culture for 3 days in conditioned medium or control medium, respectively. Immunofluorescence staining of the DNA double-strand-break marker γH2AX [32] was performed using a JBW301 mouse monoclonal anti-γH2AX antibody (Merck, Darmstadt, Germany) and an Alexa Fluor 488-conjugated goat anti-mouse secondary antibody (Thermo Fisher) [33, 34]. At least 50 nuclei were evaluated in each analysis. Cytogenetic analysis of G-banded chromosomes was performed according to standard procedures [35]. At least 25 metaphases were analyzed in each sample according to ISCN 2016 [36]. Cell viability was assessed using the CellTiter-Glo luminescent cell viability assay (Promega, Fitchburg, US) according to the manufacturer’s instructions. Luminescence was measured using a microplate reader (Tecan, Männedorf, Switzerland). ROS were analyzed using the ROS Detection Kit (PromoCell, Heidelberg, Germany) according to the manufacturer’s instructions. Luminescence was measured using a microplate reader (Tecan).

### Protein quantitation using mass spectrometry

A proteomics approach for label-free quantitation using nanoscale liquid chromatography coupled to tandem mass spectrometry (nano LC-MS/MS) was applied for comparison of proteome differences.

### Sample preparation for proteome analysis

Samples were prepared from 2 Gy irradiated MSC 4 h after irradiation and from nonirradiated control MSC. All MSC of 80% confluent T175 flasks were collected and washed three times in PBS. Afterwards, MSC were lysed in 200 μl RIPA buffer supplemented with Halt Protease Inhibitor Cocktail (100X) (Thermo Fisher) on ice for 30 min. Further, MSC conditioned medium and control medium were prepared using serum-free StemMACS MSC Expansion Media XF as stated before. Finally, samples from CD34^+^ cells were prepared at day 6 after culture for 3 days in untreated medium followed by culture for 3 days in conditioned medium or control medium, respectively. After washing the samples three times in PBS, 1×10^6^ CD34^+^ cells of each sample were lysed in 200 μl RIPA buffer supplemented with Halt Protease Inhibitor Cocktail (100X) on ice for 30 min. Lysates were stored at – 20 °C.

### Sample fractionation by SDS-PAGE and in-gel digestion

Cell culture supernatants were concentrated tenfold before SDS polyacrylamide gel electrophoresis (SDS-PAGE) by ultrafiltration (MWCO 5 kDa). Samples were heated to 95 °C for 5 min and cooled on ice prior to loading on NuPAGE 4-12% Bis-Tris gels (Thermo Fisher). SDS-PAGE was performed of all compared samples in parallel according to the manufacturer’s specification. Proteins were fixed within the polyacrylamide matrix by incubating the entire gel in 5% acetic acid in 1:1 (vol/vol) water:methanol for 30 min. After Coomassie staining (60 min) the gel slab was rinsed with water for 60 min. Each lane was excised and subdivided in three fractions according to protein complexity over standardized molecular weight ranges. Gel fractions were cut into small pieces. Subsequently, proteins were destained by 100 mM ammonium bicarbonate/acetonitrile 1:1 (vol/vol) before reduction for 30 min in 10 mM DTT and alkylation for 30 min in 50 mM iodoacetamide. Finally, proteins were digested by trypsin overnight at 37 °C. Peptides were collected from supernatant and extracted additionally from gel pieces by 1.5% formic acid in 66% acetonitrile for 15 min. Peptides from both steps were combined and dried down in a vacuum centrifuge.

### Mass spectrometry

Fractions of dried peptides were re-dissolved in 35 μl 0.1% trifluoroacetic acid and analyzed individually. For this, peptides were loaded on a 75 μm x 2 cm Acclaim C18 precolumn (Thermo Fisher) using an RSLCnano HPLC system (Thermo Fisher). Then, peptides were eluted with an aqueous-organic gradient (4-44% acetonitrile, 0.1% formic acid) for 130 min and separated on a 75 μm x 15 cm Acclaim C18 column (Thermo Fisher) with a flow rate of 300 nl/min. A Triversa Automate (Advion, Ithaca, US) was used as ion source to produce a stable electrospray, which was analyzed on a LTQ Orbitrap XL mass spectrometer (Thermo Fisher). Each scan cycle consisted of one FTMS full scan and up to 10 ITMS dependent MS/MS scans of the ten most intense ions with dynamic exclusion set to 30 sec. Mass width was set to 10 ppm and monoisotopic precursor selection was enabled. All analyses were performed in positive ion mode.

### Comparative proteome analysis

Differences in proteomes between treatment groups were analyzed by Proteome Discoverer version 2.4 (Thermo Fisher). Comparisons were made between matching sample types and fractions. CD34+ cells and MSC analyses were based on 5 replicates. For the comparison of protein supplement-free cell culture supernatants, 4 replicates were utilized. The analyses were based on at least 10 ppm mass accuracy and 1% false discovery rate. Peptides were identified using the SEQUEST algorithm and a human proteome database retrieved from UniProt (Aug. 2019, https://www.uniprot.org). Protein abundance was calculated based on intensities of unique precursor ions and limited to unmodified peptides with high confidence. Precursor ion intensities were normalized to the total peptide amount in each sample. Protein abundance ratios derived from irradiated vs. non-irradiated cell samples were calculated as median of pairwise precursor comparison of replicates to reflect the pairwise experimental design of treatments. Missing intensities were imputed based on replicates, and statistics were calculated by background based ANOVA. In cell culture supernatants, the number of required background elements was insufficient for background based ANOVA. Therefore, t-tests were calculated for individual proteins. Furthermore, differences in cell culture supernatant were based on the top three scored unique peptides to account for protein processing, such as signal peptide truncation, etc.. All protein identifications were filtered for a required minimum of at least two unique peptides. A minimum of two distinct peptides with similar regulation was utilized as a requirement for calculated ratios during manual inspection. In addition, a minimum detection in at least three replicates was an essential inclusion criterion for calculated ratios during manual inspection. Tables summarizing differences in proteomes between treatment groups meet all criteria described above and include corresponding *p* values.

### Statistical analysis

Proteomic data were analyzed as outlined in the section above. All other statistical calculations were done with SAS software, release 9.4 (SAS Institute, Cary, US). For comparisons between treated groups and controls, Wilcoxon two-sample tests were used. One sample t-tests were used in order to investigate if mean fold changes (fc) were different from 1.

## Results

### Validation of cell-free MSC conditioned medium

To ensure that MSC conditioned medium was cell-free in our experiments only centrifuged supernatants of MSC conditioned medium were used. In addition, (a) microscopic evaluation of supernatants in a Neubauer counting chamber, (b) control experiments (n = 3) with sterile filtered supernatants and (c) cytogenetic cross-over experiments (n = 2) using sexually divergent CD34+ cells and MSC were performed, which demonstrated no transfer of MSC in our experiments.

### ROS in MSC and CD34+ cells

ROS were analyzed in 2 Gy irradiated MSC samples (n = 8) at 4 h after irradiation and in non-irradiated control MSC. Increased ROS levels (*p* = 0.0105) were detected in irradiated MSC (fc =1.8 ± 0.2; mean ± standard error of mean (SEM)) when compared to nonirradiated MSC (fc = 1) (Fig. 1a). Furthermore, ROS were analyzed in CD34+ cell samples (n = 5) expanded for 3 days in untreated medium followed by culture for 3 days in conditioned medium or control medium, respectively. ROS levels tended to be increased (*p* = 0.2206) in CD34+ cells grown in conditioned medium (fc = 1.2 ± 0.2) when compared to ROS levels in CD34+ cells grown in control medium (fc = 1) (Fig. 1b).

**Fig. 1.**
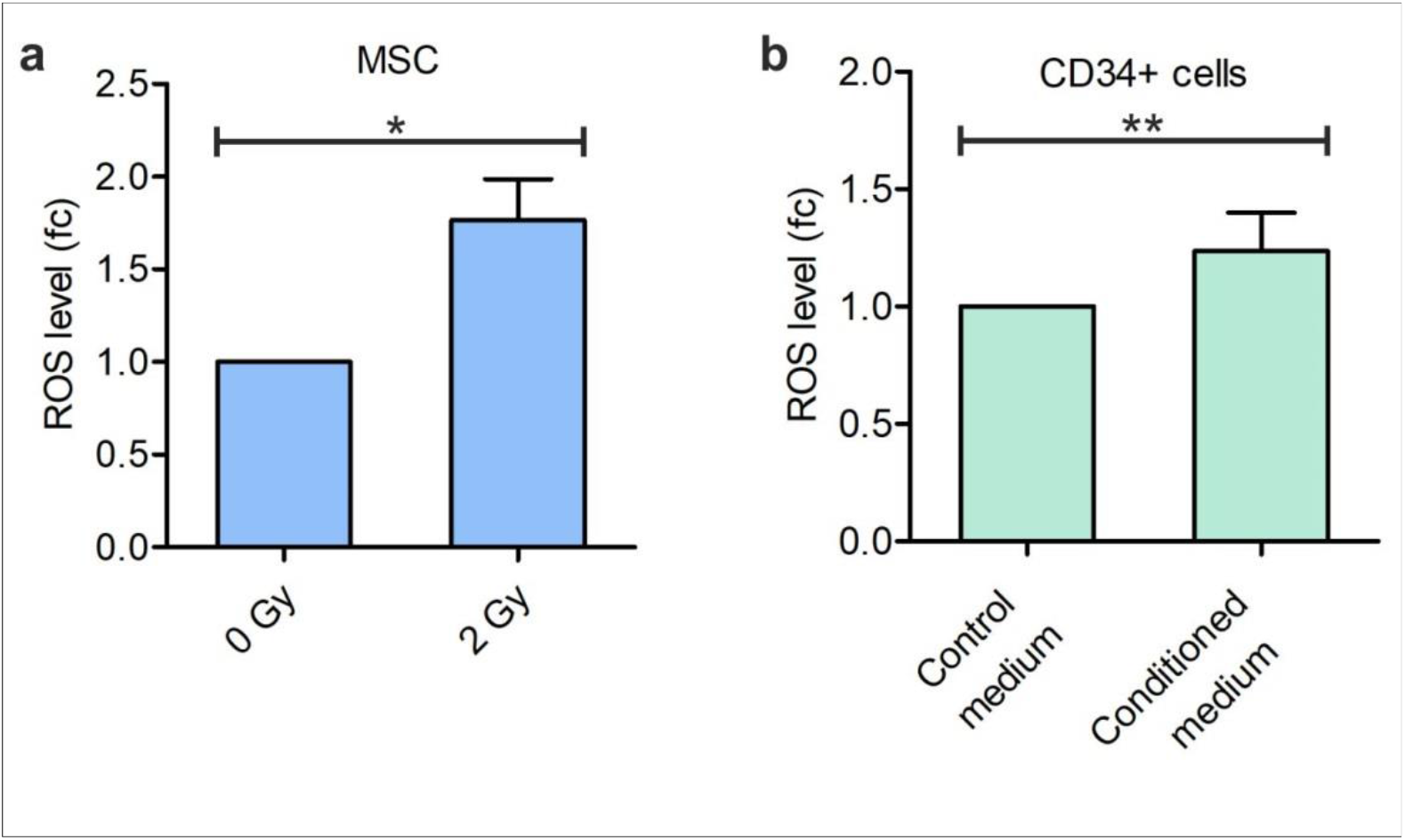
Reactive oxygen species (ROS) levels in irradiated mesenchymal stromal cells (MSC) and CD34+ cells grown in MSC conditioned medium. **a** ROS levels in 2 Gy irradiated MSC at 4 h after irradiation. \**p* = 0.0105. **b** ROS levels in CD34+ cells grown for 3 days in medium conditioned by 2 Gy irradiated MSC. \*\**p* = 0.2206. *fc* fold change.

### DNA damage in CD34+ cells

γH2AX foci were analyzed in CD34+ cell samples (n = 9) expanded for 3 days in untreated medium followed by culture for 3 days in conditioned medium or control medium, respectively. γH2AX foci levels were increased (*p* = 0.0003) in CD34+ cells grown in conditioned medium (2.8 ± 0.5 γH2AX foci per CD34+ cell; mean ± SEM) when compared to γH2AX foci levels in CD34+ cells grown in control medium (0.5 ± 0.1 γH2AX foci per CD34+ cell) (Fig. 2a, b).

**Fig. 2.**
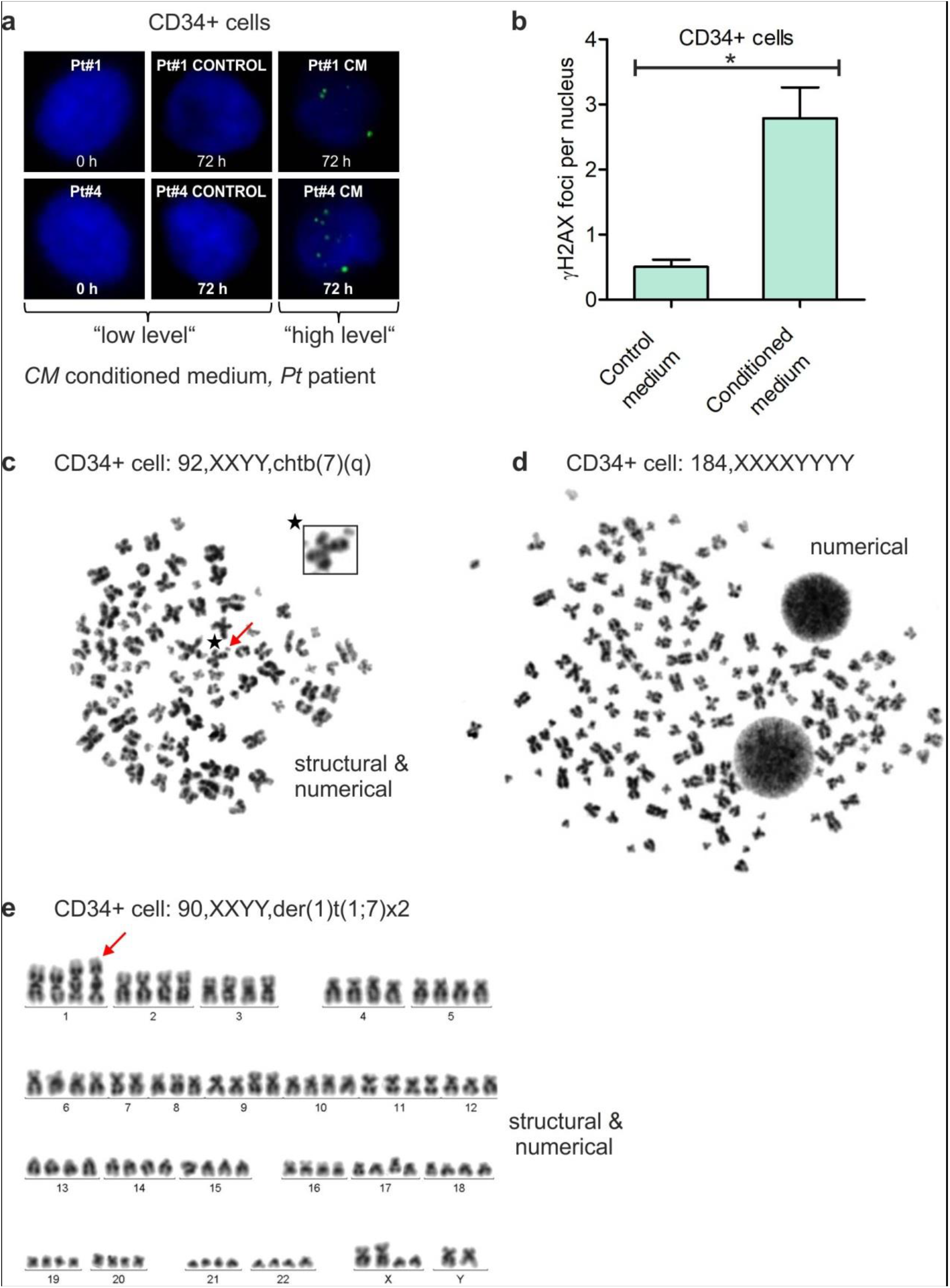
Radiation-induced bystander effects in CD34+ cells. **a** Exemplary immunofluorescence images of γH2AX foci (green, Alexa 488) in nuclei (blue, DAPI) of CD34+ cells grown for 3 days in medium conditioned by 2 Gy irradiated mesenchymal stromal cells (MSC). **b** The numbers of γH2AX foci were increased in CD34+ cells grown for 3 days in medium conditioned by 2 Gy irradiated MSC when compared to the numbers of γH2AX foci in CD34+ cells grown in control medium. \**p* = 0.0003. **c-d** Exemplary aberrant metaphases of different donor CD34+ cells grown for 3 days in MSC conditioned medium. **e** Exemplary aberrant karyotype of a donor CD34+ cell grown for 3 days in MSC conditioned medium.

### Chromosomal instability in CD34+ cells

Metaphases were analyzed in CD34+ cell samples (n = 12) expanded for 3 days in untreated medium followed by culture for 3 days in conditioned medium or control medium, respectively (Fig. 2c-e, Table 1). Structural and numerical chromosomal aberrations were detected in 50% and 92% of CD34+ cell samples grown in MSC conditioned medium, respectively, when compared to normal karyotypes detected in CD34+ cell samples grown in control medium. In particular, chromatid breaks (chtb), e.g., chtb(5q), chtb(6p), chtb(7q), chtb(10q), chtb(11q) and chtb(13q), translocations, e.g., der(1)t(1;7) and aneuploidies, e.g., tetraploidies and octoploidies, were observed in CD34+ cells grown in conditioned medium. Finally, the estimated mitotic rates as determined by the number of dividing cells among all cells were similar in CD34+ cells grown in conditioned medium and in CD34+ cells grown in control medium.

### Viability of CD34+ cells

Viability was assessed in CD34+ cell samples (n = 10) grown for 3 days in untreated medium followed by culture for 3 days in conditioned medium or control medium, respectively. Viability of CD34+ cells grown in conditioned medium (fc = 1.1 ± 0.1; mean ± SEM) was similar when compared to viability of CD34+ cells grown in control medium (fc = 1; data not shown).

### Proteome analysis in MSC, MSC conditioned medium and CD34+ cells

Comparative proteome analysis was performed in patient samples with (a) lysates of irradiated and non-irradiated MSC, (b) MSC conditioned and control medium and (c) lysates of CD34+ cells grown in conditioned and control medium (Figs. 3, 4, Table 2). In MSC, 31 of 1924 identified proteins (1.6%) were regulated at least twofold within 4 hours upon a single irradiation dose of 2 Gy compared to controls. The majority was upregulated (94%) and about half participate in translation, protein folding as well as protein degradation. Six altered proteins are part of the cytoskeleton and participate in its dynamic regulation. Four are members of nuclear transport mechanisms and the nuclear pore complex. The remaining participate in energy metabolism (e.g., glycolysis), oxidative stress detoxification, cell-cell/matrix interaction and transmembrane signaling.

**Fig. 3.**
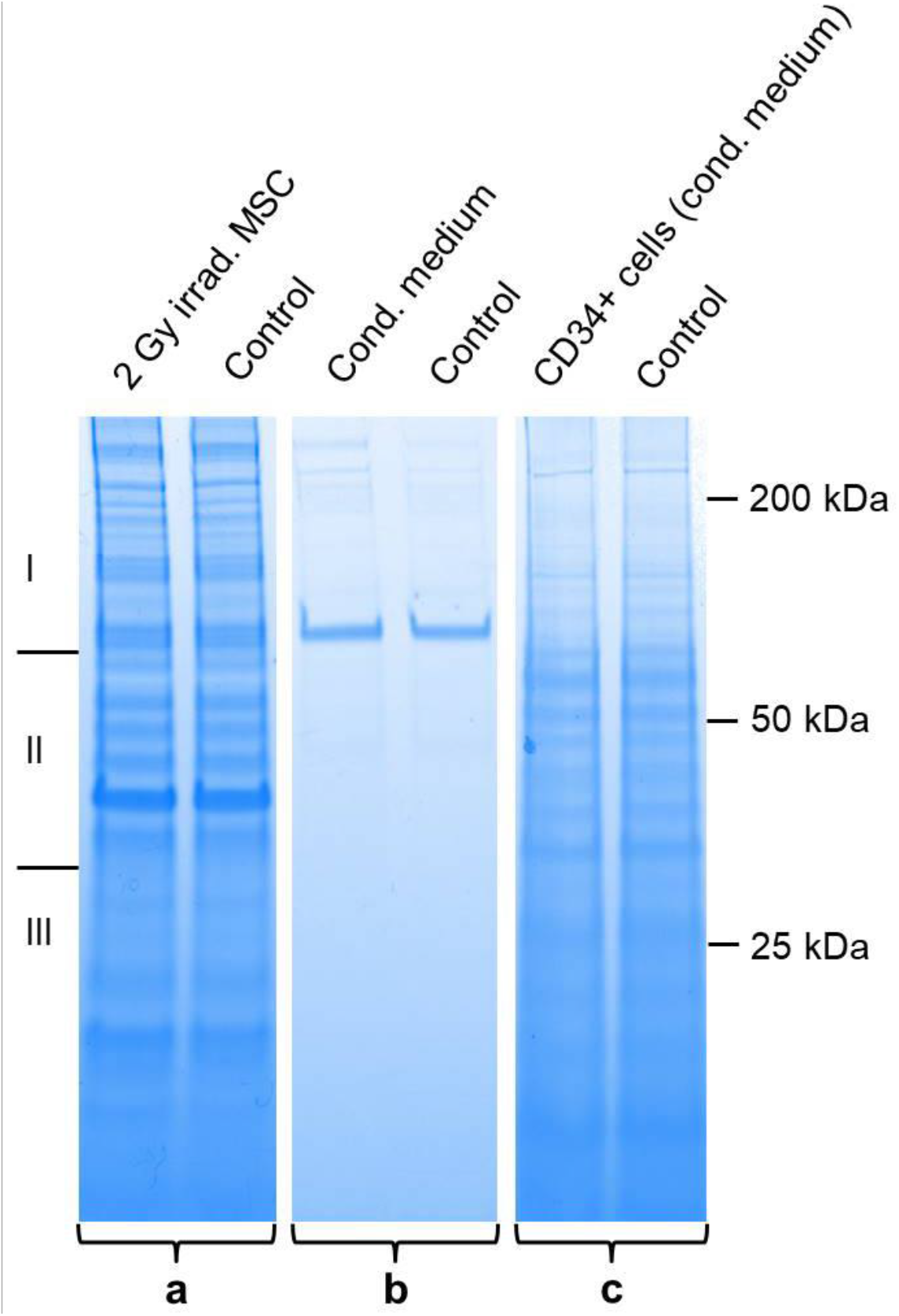
SDS-PAGE. **a** Lysates of irradiated and non-irradiated MSC, **b** MSC conditioned and control medium and **c** lysates of CD34+ cells grown in conditioned and control medium.

**Fig. 4.**
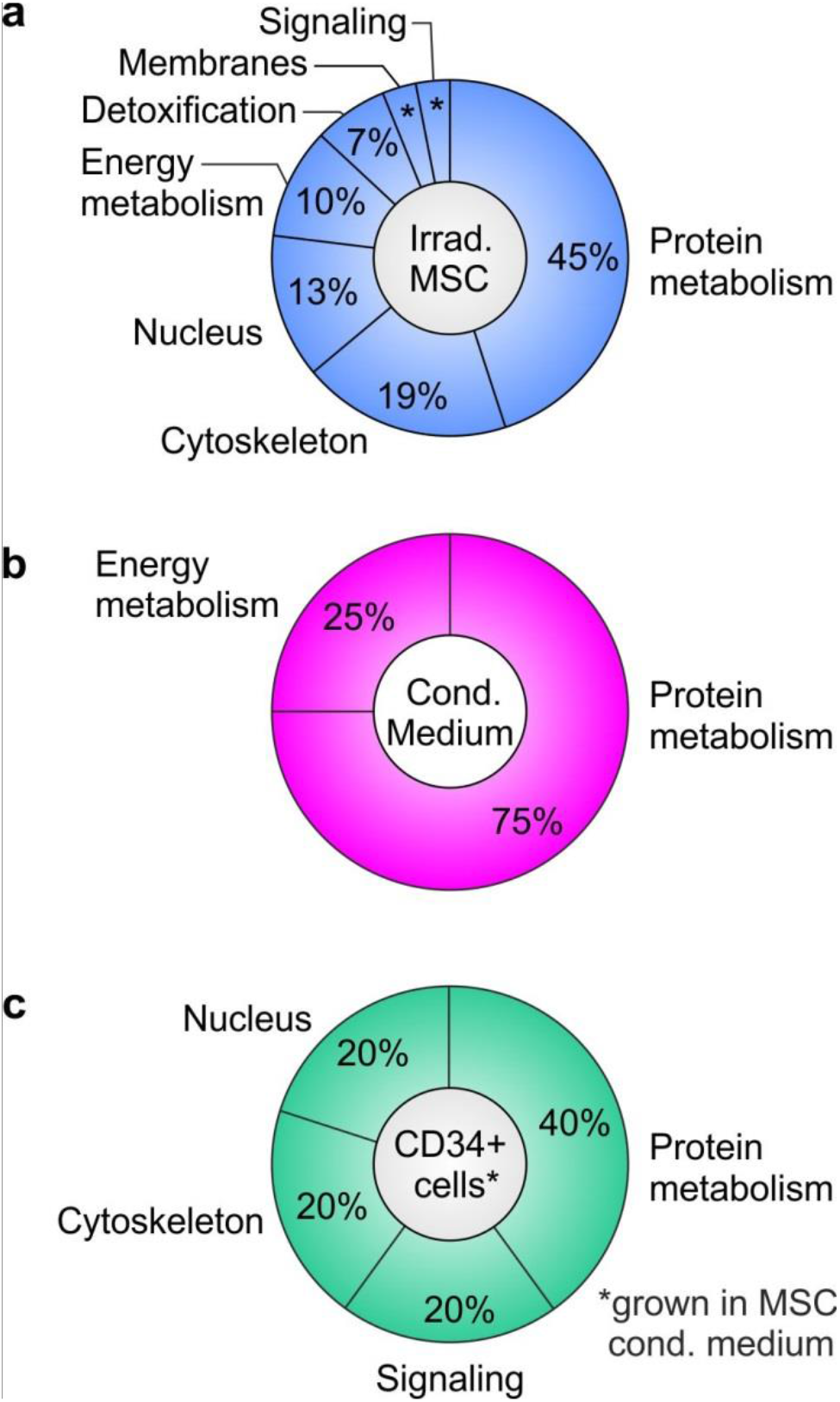
Proteome alterations according to categories in mesenchymal stromal cells (MSC), MSC conditioned medium and CD34+ cells. **a** Proteome shifts in irradiated MSC. * 3%. **b** Proteome shifts in MSC conditioned medium. **c** Proteome shifts in CD34+ cells grown for 3 days in MSC conditioned medium.

**Table 2.**
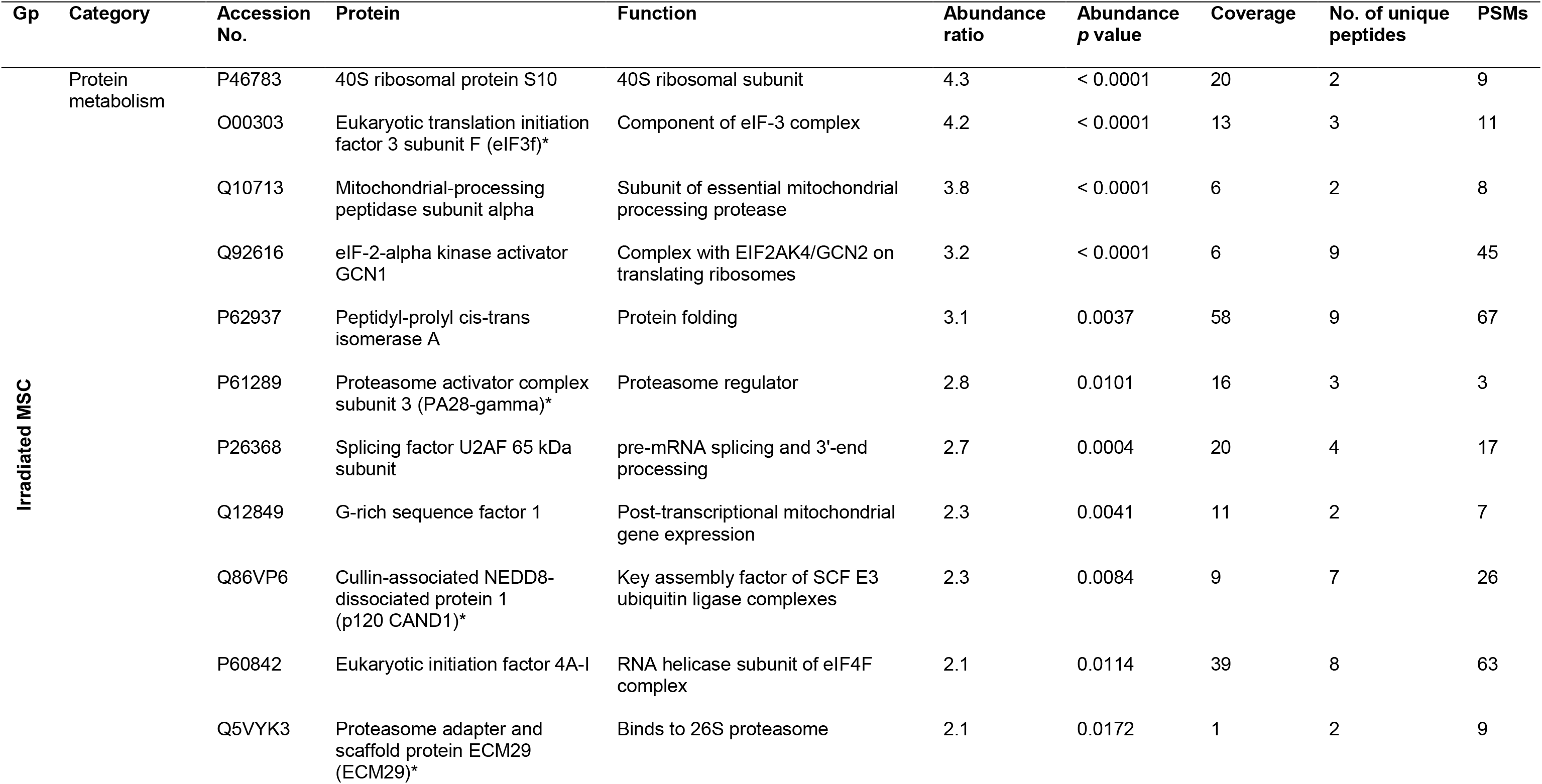
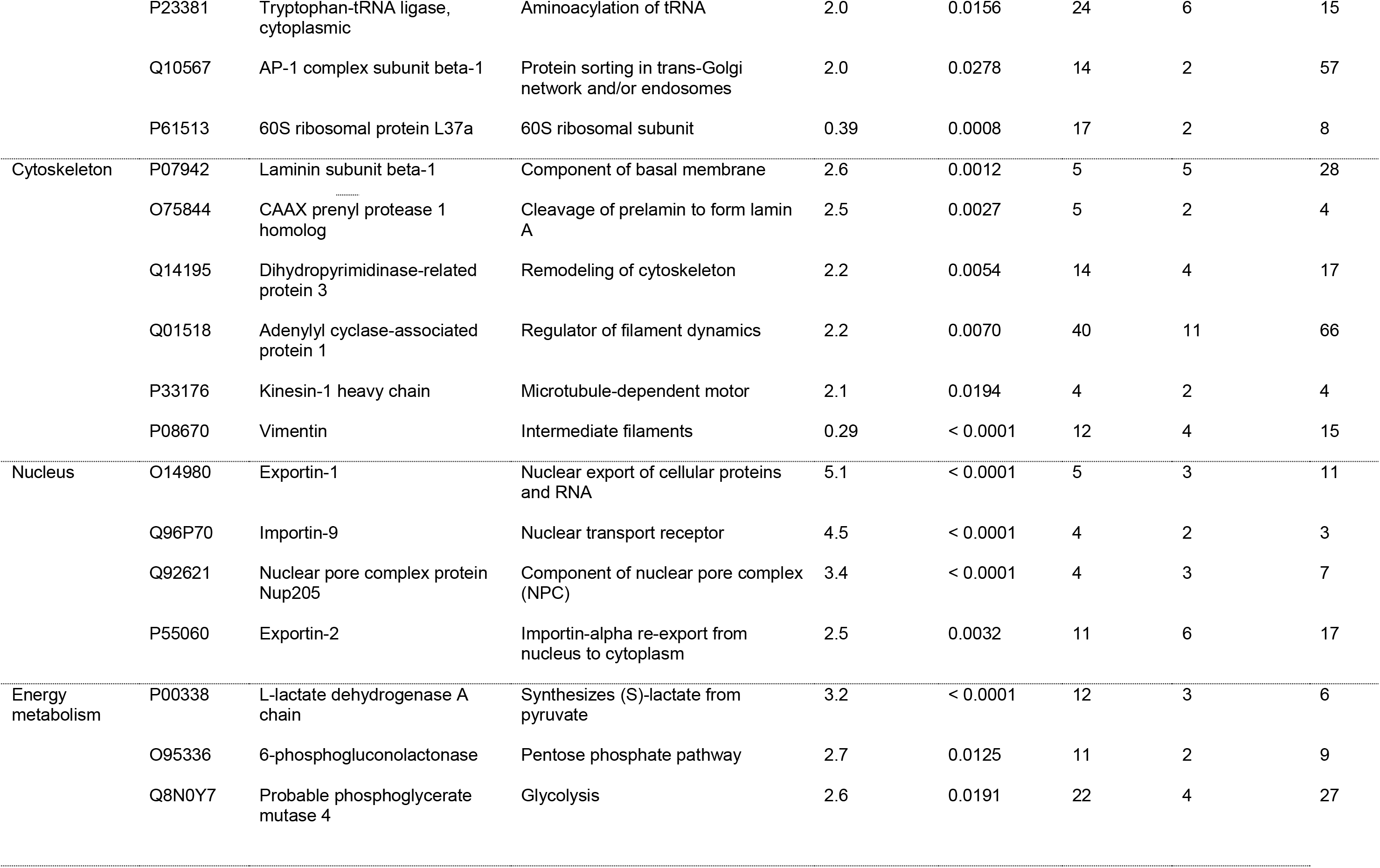
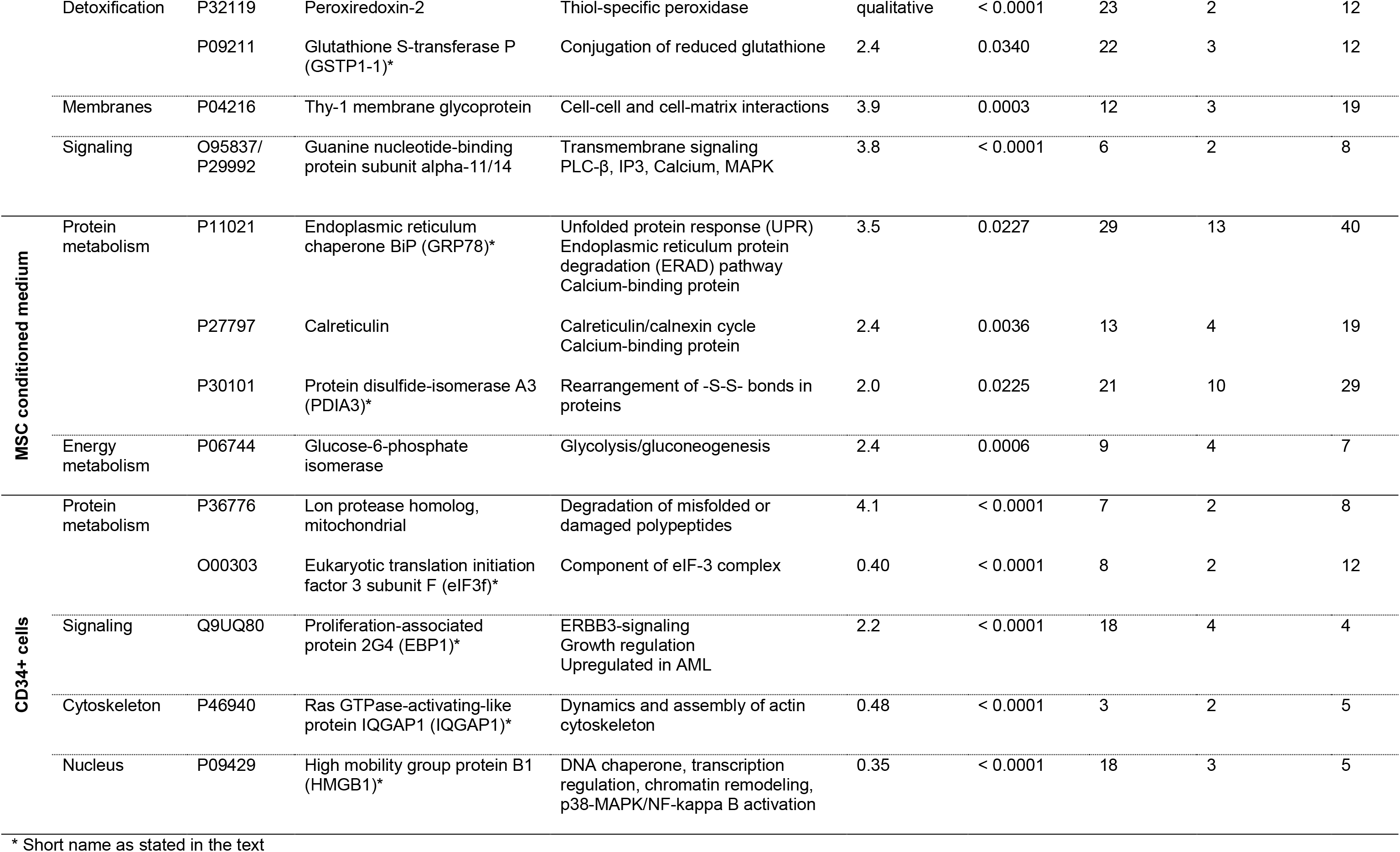
Proteome alterations in 2 Gy irradiated mesenchymal stromal cells, MSC conditioned medium and CD34+ cells grown in MSC conditioned medium in comparison to controls. *Gp* group; *PSMs* peptide-to-spectrum matches.

In the corresponding conditioned medium of MSC, 4 of 265 identified proteins (1.5%) were found increased in their abundance by factor 2 or higher 4 h after irradiation vs. controls. Remarkably, three of these proteins are key proteins in the endoplasmatic reticulum (ER) and known for their role in protein folding as well as protein quality control. Besides one upregulated member of the glycolysis, no other proteins were differentially abundant in conditioned medium.

Exposure of CD34+ cells to the conditioned medium of irradiated MSC for three days induced quantitative changes of a minimum factor 2 in 5 of 2003 identified proteins (0.25%). Similar to MSC, affected proteins participate in translation, protein degradation and cytoskeleton dynamics. Notably, eIF3f was lower abundant in CD34+ cells, whereas it was higher abundant in MSC in conjunction with irradiation. Unique to the response in CD34+ cells were changes in proteins participating in transcriptional regulation/chromatin remodeling and ERBB3 regulation. Overall, the response in CD34+ cells to conditioned medium affected much less proteins than in MSC, which were directly exposed to irradiation.

## Discussion

The aim of our study was to analyze genetic alterations induced by DNA damage signaling from irradiated MSC to human CD34+ cells as a potential mechanism of MN initiation. For this purpose, RIBE were analyzed in human CD34+ cells grown in medium conditioned by 2 Gy irradiated human MSC. Notably, increased numbers of γH2AX foci as well as structural and numerical chromosomal aberrations were detected in CD34+ cells grown in MSC conditioned medium when compared to CD34+ cells grown in control medium. The increased numbers of γH2AX foci in CD34+ cells grown in MSC conditioned medium may not only indicate critical DNA damage potentially contributing to MN initiation, e.g., by activation of oncogenes or inactivation of tumor suppressor genes [37]. In addition, γH2AX foci may indicate DSB [32] involved in chromosomal rearrangements such as deletions, inversions and translocations. Indeed, t-MN related chromosomal aberrations were found in CD34+ cells grown in MSC conditioned medium when compared to whole chromosomes in CD34+ cells grown in control medium. Particularly, chtb(5q), chtb(7q), chtb(11q) and chtb(13q), that were found in CD34+ cells grown in MSC conditioned medium, coincided well with del(5q), del(7q), t(11q23.3) and del(13q), that are present in about 42%, 49%, 3% and < 5% of t-MN, respectively [1, 6]. In addition, t-MN related aneuploidies, e.g., tetraploidies and octoploidies, were detected in CD34+ cells grown in MSC conditioned medium. Numerical chromosomal aberrations are caused by defects in mitosis, e.g., chromosomal non-disjunction and cytokinesis failure [38]. Moreover, tetraploidies are hallmark precursor lesions in diverse cancers, e.g., cervical cancer and neuroblastoma and occur in about 1% of AML but 13% of t-AML cases [38, 39]. As tetraploid cells harbor 4n centrosomes, multipolar spindles may form potentially driving a CIN phenotype. With ongoing dedifferentiation of CD34+ cells CIN may further aggravate in the course of disease evolution, e.g., by frequent inactivation of *TP53*, which may result in rapid t-MN development [38]. Finally, the increased numbers of γH2AX foci and chromosomal aberrations did not seem to affect overall viability of CD34+ cells within the observation period as viability was similar in CD34+ cells grown in conditioned medium and in CD34+ cells grown in control medium.

ROS were analyzed in irradiated MSC and CD34+ cells grown in MSC conditioned medium for their potential contribution to bystander signaling from irradiated MSC to CD34+ cells. Increased ROS levels were detected in irradiated MSC and in CD34+ cells grown in MSC conditioned medium. While ROS are known genotoxic molecules generated by endogenous and exogenous sources in each cell, ROS may also function as important regulators of intracellular signaling pathways, e.g., by covalent modification of specific cysteine residues in redox-sensitive target proteins [40]. Oxidation of specific cysteine residues in turn can lead to reversible modification of enzyme activity [40] with effects on diverse pathways including metabolism, differentiation and proliferation [41]. Hence, ROS may not only induce DNA damage but also dysregulate cellular pathways, thereby contributing to oncogenic transformation of CD34+ cells.

In order to identify potential mediators for the observed oncogenic transformation in CD34+ cells as well as mechanisms leading to their release in MSC and transduction in CD34+ cells, comparative proteome analyses were performed in three tiers of (a) irradiated MSC, (b) MSC conditioned medium and (c) CD34+ cells grown in MSC conditioned medium. Among these three comparisons, irradiated MSC showed the largest change in proteome, which is in accordance with the impact of the primary stimulus. Still, the response can be regarded as rather moderate, because only 1.6% of the analyzed proteome was altered by a factor 2 or higher. An underlying mechanism might be the relative radioresistance of MSC [42]. The majority of altered proteins in response to irradiation take part in translation, protein folding as well as protein degradation, indicating disturbed protein homeostasis and required replacement, repair and degradation of proteins. Interestingly, three of the few quantitatively altered proteins in MSC conditioned medium upon irradiation were key ER proteins involved in protein folding and their quality control. The highest increase was observed for GRP78, an ER chaperone, which dissociates from luminal domains of IRE1, PERK and ATF6 in consequence of ER stress resulting in activation of the unfolded protein response (UPR) [43] and promotion of the ER-associated protein degradation pathway (ERAD) [44]. In turn, ERAD relies on substrate degradation via the ubiquitin-proteasome system. Notably, two proteasome activator proteins (ECM29 and PA28-gamma) as well as a key assembly factor of SCF E3 ubiquitin ligase complexes (p120 CAND1) were all increased in irradiated MSC. Altogether the results indicate that irradiation resulted in ER stress. The stress response may be induced in part by associated ROS. At proteome level, MSC responded to increased oxidative stress by elevating levels of peroxiredoxin-2 and GSTP1-1.

The perception about GRP78 has changed over the past decade, as a growing number of signaling processes become apparent, which are not related to its canonical role in the ER [45, 46]. It appears that GRP78 is not exclusively present in the ER but can be relocated to the cell surface (csGRP78) or even secreted into the extracellular medium (sGRP78). Both have been described to confer critical roles in the context of cancer development and cell survival [45, 46]. For example, sGRP78 can act as a pro-apoptotic ligand of csGRP78 on pancreatic β-cells [47], but as a mediator of pro-survival kinase signaling in endothelial cells [48]. In addition, csGRP78 plays a mechanistic role in PI3K/AKT driven leukemogenesis [49] and in Cripto/csGRP78 regulated hematopoietic stem cell survival [50]. Therefore, monitoring of sGRP78 and targeting of csGRP78 is evaluated in anti-cancer therapy [45, 46]. Considering these emerging roles of GRP78, non-canonical csGRP78 signaling may impact the survival of CD34+ cells harboring genetic aberrations and contribute to oncogenic bystander signaling. The remaining two ER proteins with increased abundance upon irradiation in MSC conditioned medium were PDIA3 and calreticulin. PDIA3 catalyzes the rearrangement of disulfide bonds [51] and thereby enables correct folding of newly-synthesized glyco-proteins [52]. In addition, it interacts with the ER resident calcium binding lectins calreticulin and calnexin. Similar to the function of proteins altered in response to irradiation in MSC, calreticulin and calnexin participate in protein quality control and folding, more specifically, in a process known as the calreticulin/calnexin cycle [53]. The fact that three ER proteins with related function were specifically increased in the conditioned medium upon MSC irradiation, while the vast majority of other cytosolic and ER proteins were unaffected, suggests a specific release rather than uncontrolled cell lysis or unspecific cellular loss of the ER.

In CD34+ cells, the conditioned medium from irradiated MSC induced only minute detectable changes at proteome level after three days of exposure. Individual proteins participating in degradation, translation and cytoskeleton dynamics represent similar processes affected in MSC. Unique to CD34+ cells were proteins participating in chromatin remodeling (HMGB1) and ERBB3 signaling (EBP1). In particular EBP1 has oncogenic potential [54] and is highly expressed in AML cells [55], but HMGB1 assumes a number of roles in cancer development as well [56]. In addition, IQGAP1 can promote malign development [57]. As a consequence, several modes of action, which work individually or in conjunction, may transduce radiation-induced bystander signaling in effector cells.

Our data describe a sequence of cellular events from the primary response of irradiated MSC, over transmission of genotoxic signals in conditioned medium to the induction of mechanisms leading to critical DNA damage and CIN in CD34+ cells (Fig. 5). Ultimately, such genetic aberrations in effector cells have the potential for MN development. The results provide a fundamental basis for in-depth mechanistic research and novel therapeutic interventions to reduce the risk of t-MN development. Accordingly, antioxidants, such as N-acetylcysteine, might be able to counteract ROS in MSC and HSPC, thereby diminishing the risk for t-MN after irradiation. In addition, monoclonal antibodies (e.g., MAb159) [58] and peptidomimetics (e.g., BMTP-78) [59] targeting non-canonical csGRP78 signaling hold the potential to reduce the risk of t-MN.

**Fig. 5.**
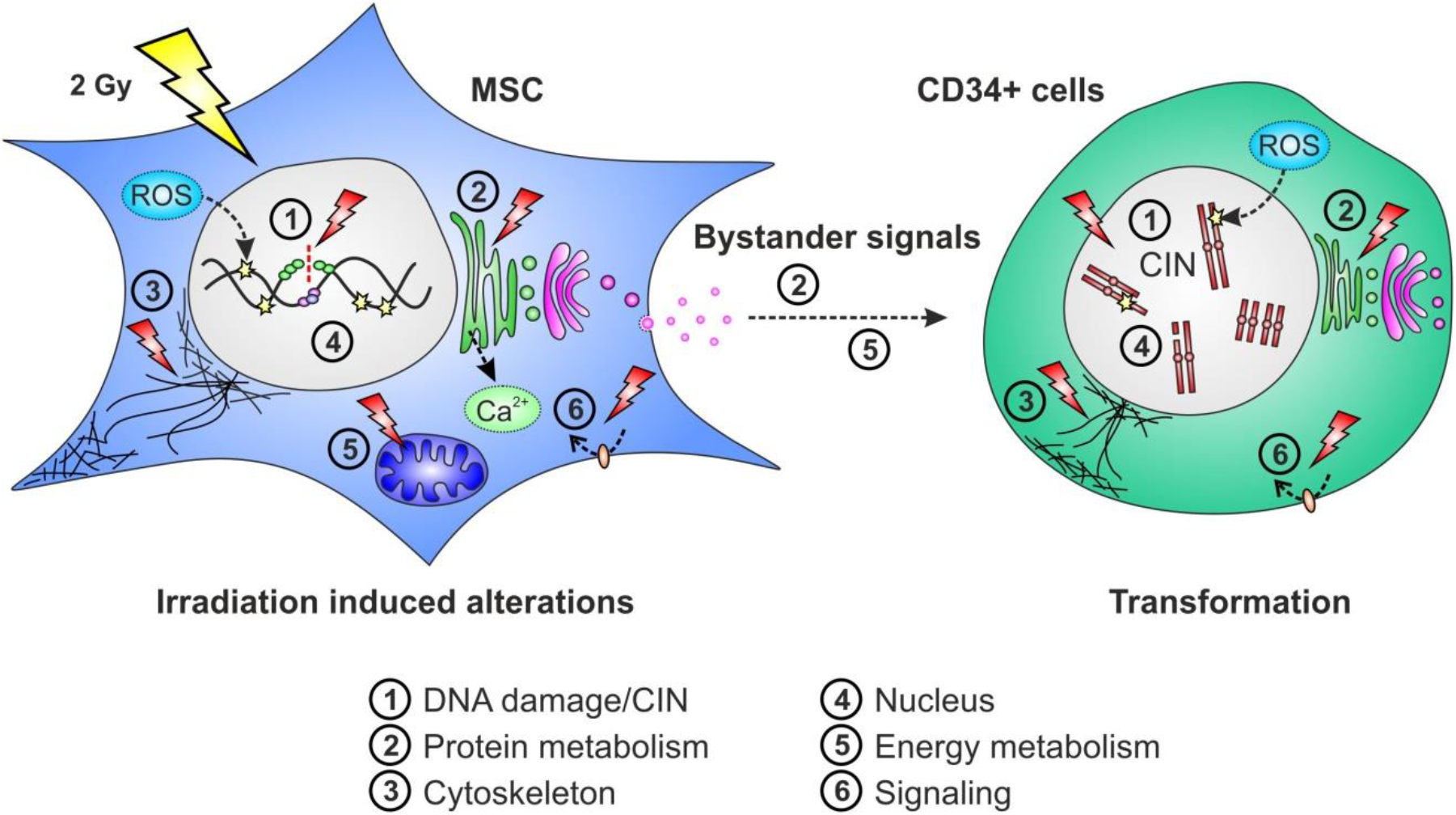
Model of bystander signaling in mesenchymal stromal cells (MSC) and CD34+ cells. Irradiation of MSC may induce DNA damage (**1**) directly and indirectly by reactive oxygen species (ROS). Induced protein shifts in MSC may interfere with protein metabolism (**2**), cytoskeleton (**3**), nucleus (**4**), energy metabolism (**5**) and signaling (**6**). Genotoxic bystander signals (**2**) and (**5**) may be transmitted to CD34+ cells. In CD34+ cells the signals may induce ROS and protein shifts interfering with protein metabolism (**2**), cytoskeleton (**3**), nucleus (**4**) and signaling (**6**) ultimately leading to DNA damage and CIN (**1**).

In conclusion, genotoxic signaling by irradiated MSC emerges as a major pathomechanism in the initiation of certain MN offering the opportunity to take advantage of targeted therapeutic interventions. Specifically, our data suggest that bystander signals released by irradiated MSC, such as GRP78, are potential mediators of DNA damage and CIN in CD34+ cells, thereby providing a strong mechanistic link to the initiation of MN. More work is necessary to dissect the signaling pathways behind such oncogenic mediators which may define the targets of next-generation anti-leukemic drugs.

## Data availability

Primary proteomic data are available from the corresponding author upon request.

## Acknowledgements

HDP and AF received grants from Deutsche José Carreras Leukämie-Stiftung (DJCLS 14 R/2017). DN is an endowed Professor of the same foundation (DJCLS H 03/01).

## Author contributions

VK performed RIBE analysis, OD contributed proteomic expertise, VC conducted mass spectrometry, MB and HK irradiated MSC, AJ and HR supported femoral head collection, CW performed statistical analysis, SB supported RIBE analysis, JF, BS, WS, DN and WKH provided expertise, AF contributed cytogenetic expertise, HDP designed the study and wrote the manuscript.

## 1. Introduction

Genotoxic bystander signals released from irradiated human mesenchymal stromal cells (MSC) may induce radiation-induced bystander effects (RIBE) in non-irradiated human hematopoietic stem and progenitor cells (HSPC) potentially initiating myeloid neoplasms (MN). In the 2016 WHO classification, MN that arise after irradiation therapy are referred to as therapy-related MN (t-MN) [1]. As t-MN are characterized by high-risk genetic alterations [2, 3] and a particularly worse prognosis [4, 5], novel therapeutic strategies are urgently needed.

Generally, RIBE describe ‘out-of-field’ effects of irradiation in non-irradiated cells that are comparable to effects in irradiated cells. RIBE may emerge as DNA damage (e.g., increased γH2AX foci, gene mutations, chromosomal aberrations, micronuclei), cell death (e.g., apoptosis, necrosis) and induction of cell survival mechanisms (e.g., adaptive response, DNA repair) [6–9]. Bystander signals are assumed to be initiated in irradiated cells by calcium fluxes [10] and mitochondrial metabolites [11–13]. Then, small molecules like nitric oxide (NO) [14], reactive oxygen species (ROS) [15], nuclear factor-kappa B (NF-kappa B) [13], and transforming growth factor beta-1 (TGFbeta-1) [16, 17] may pass through cell membranes and gap junctions from the intracellular towards the extracellular space [18, 19]. Hereupon, the bystander signals might be transmitted to non-irradiated cells that are referred to as bystander cells. Finally, ROS generated by NADH oxidases [20] and distinct RIBE mediators may be induced in affected bystander cells, thereby potentially initiating malignant transformation.

The analysis of bystander signals is a cutting-edge field in leukemia research. Here, irradiated healthy human MSC and healthy human CD34+ cells from the same donors were investigated in an *in vitro* model system that enables characterization of genotoxic signaling factors. Specifically, molecular size fractions of MSC conditioned medium of approximate (I) < 10 kDa, (II) 10 – 100 kDa and (III) > 100 kDa molecular weight were used for culture of CD34+ cells of the same donors. Afterwards, RIBE were analyzed in exposed CD34+ cells in terms of DNA damage and chromosomal instability (CIN). The data may provide important information on the fraction of interest in MSC conditioned medium to be analyzed most profitable by in-depth proteome analysis for the identification of key bystander signals, which might contribute to the development of next-generation anti-leukemic drugs.

## 2. Experiments

### 2.1. Preparation of human femoral heads

This study was approved by the Ethics Committee II, Medical Faculty Mannheim, Heidelberg University (no. 2019-1128N). Procedures were performed in accordance with the local ethical standards and the principles of the 1964 Helsinki Declaration and its later amendments. Written informed consent was obtained from all study participants. Femoral heads were collected from 6 patients with coxarthrosis (1 female, 5 males, mean age: 68 years) undergoing hip replacement.

### 2.2. Isolation of human MSC

Bones were broken into fragments and incubated for 1 hour at 37 °C in phosphate-buffered saline (PBS) supplemented with 1 mg/ml collagenase type I (Thermo Fisher, Waltham, US). Supernatants were filtered in a cell strainer with 100 μm nylon mesh pores (Greiner Bio-One, Kremsmünster, Austria). Afterwards, bone fragments retained in the cell strainer were transferred into StemMACS MSC Expansion Media XF (Miltenyi Biotec, Bergisch Gladbach, Germany) supplemented with 1% penicillin/streptomycin. Then, adherent MSC were expanded in T175 flasks in a humidified 5% CO_2_ atmosphere at 37 °C and passaged at 80% confluency.

### 2.3. Isolation of human CD34+ cells

CD34+ cells were isolated from bone marrow mononuclear cells by Ficoll density gradient centrifugation and magnetic-activated cell sorting using CD34 antibody-conjugated microbeads (Miltenyi Biotec). CD34+ cells were grown in StemSpan SFEM II medium (Stemcell Technologies, Vancouver, Canada) supplemented with StemSpan Myeloid Expansion supplement (SCF, TPO, G-CSF, GM-CSF) (Stemcell Technologies) and 1% penicillin/streptomycin in a humidified 5% CO_2_ atmosphere at 37 °C.

### 2.4. Preparation of fractions of MSC conditioned medium

MSC were grown in T175 flasks until reaching 80% confluency. MSC were rinsed in PBS and fresh StemSpan SFEM II medium was added. Afterwards, MSC were 2 Gy irradiated by 6 MV x-rays in a Versa HD linear accelerator (Elekta, Stockholm, Sweden), while control MSC were not irradiated. MSC conditioned medium and control medium were obtained from irradiated and non-irradiated MSC, respectively, after 4 h incubation at 37 °C. The collected medium was centrifuged (1.200 rpm, 10 min) and supernatants were filtered through 10 kDa molecular weight cut-off (MWCO) ultrafiltration centrifugal filter units (Amicon Ultra, Merck, Darmstadt, Germany) to obtain (I) approximate < 10 kDa fractions of MSC conditioned and control medium, respectively. Next, the supernatant above the filter was adjusted with fresh medium to the original volume and filtered through 100 kDa MWCO ultrafiltration centrifugal filter units to obtain (II) approximate 10 – 100 kDa fractions of MSC conditioned and control medium, respectively. Finally, the supernatant above the filter was adjusted with fresh medium to the original volume and then contained (III) approximate > 100 kDa fractions of MSC conditioned and control medium, respectively. The distinct fractions (I) – (III) of MSC conditioned and control medium were stored at −20 °C.

### 2.5. Heat inactivation of MSC conditioned and control medium

Heat inactivation of RIBE mediators in un-/fractionated MSC conditioned medium and un-/ fractionated control medium was performed by incubation at 75 °C for 20 min.

### 2.6. RIBE analysis

RIBE were analyzed in CD34+ cell samples (6 patients) at day 6 after culture for 3 days in native medium followed by culture for 3 days in un-/fractionated MSC conditioned medium or in un-/ fractionated control medium, respectively. In addition, experiments with CD34+ cell samples (2 patients) were performed with MSC conditioned medium after heat inactivation.

### 2.7. Immunofluorescence staining of γH2AX

Immunofluorescence staining of γH2AX was performed in CD34+ cells using a JBW301 mouse monoclonal anti-γH2AX antibody (Merck) and an Alexa Fluor 488-conjugated goat anti-mouse secondary antibody (Thermo Fisher) [21, 22]. At least 50 nuclei were analyzed in each sample.

### 2.8. Cytogenetic analysis

Cytogenetic analysis of G-banded chromosomes was performed in CD34+ cells according to standard procedures [23]. At least 25 metaphases were analyzed in each sample following the ISCN 2016 [24]. Sporadic chromosomal alterations (e.g., chromatid breaks (chtb), chromosome breaks, trisomy) were included in the karyotype (non-clonal events) when detected in at least one metaphase. Because tetraploid/octaploid metaphases were detected at low frequency in CD34+ cells grown in control medium as well, they were only included in karyotypes in case of clonality (tetraploidy and/or octaploidy in two or more metaphases) according to the ISCN 2016.

### 2.9. Statistical analysis

Statistical analysis was performed with SAS software, release 9.4 (SAS Institute, Cary, US). For quantitative variables, mean values and standard deviations were calculated. Categorical factors are presented with absolute and relative frequencies. In order to compare more than two groups, Kruskal-Wallis tests were performed. For pairwise group comparisons, exact Wilcoxon two-sample tests were used. In general, test results with p < 0.05 was considered as statistically significant.

## 3. Results

### 3.1. Evaluation of cell-free MSC conditioned medium

In order to prevent a transfer of MSC by MSC conditioned medium to the CD34+ cell cultures only centrifuged supernatants were used. In addition, (i) microscopic evaluation of supernatants in a Neubauer counting chamber, (ii) sterile filtration of supernatants and (iii) cytogenetic cross-over experiments using sexually divergent CD34+ cells and MSC could exclude transfer of MSC to the CD34+ cell cultures in our experiments.

### 3.2. DNA damage in human CD34+ cells

γH2AX foci were analyzed in human CD34+ cell samples (4 patients; ∑ 32 samples) expanded for 3 days in native medium followed by culture for 3 days in un-/fractionated MSC conditioned or un-/fractionated control medium, respectively (Figure 1a). Increased numbers of γH2AX foci (general *p* = 0.0068 [Kruskal-Wallis test]; pairwise comparison each *p* = 0.0286 [Wilcoxon two-sample test]) were detected in CD34+ cells grown in the (II) 10 – 100 kDa fraction of MSC conditioned medium (0.67 ± 0.10 γH2AX foci per CD34+ cell; mean ± standard error of mean) when compared to numbers of γH2AX foci in CD34+ cells grown in (I) < 10 kDa (0.19 ± 0.01 γH2AX foci per CD34+ cell) and (III) > 100 kDa (0.23 ± 0.04 γH2AX foci per CD34+ cell) fractions or in un-/fractionated control medium (0.12 ± 0.01 γH2AX foci per CD34+ cell). Since γH2AX foci are a marker of DNA double-strand breaks (DSB), our findings suggest that DNA damage signaling factors mainly localize in the (II) 10 – 100 kDa fraction of MSC conditioned medium.

**Figure 1.**
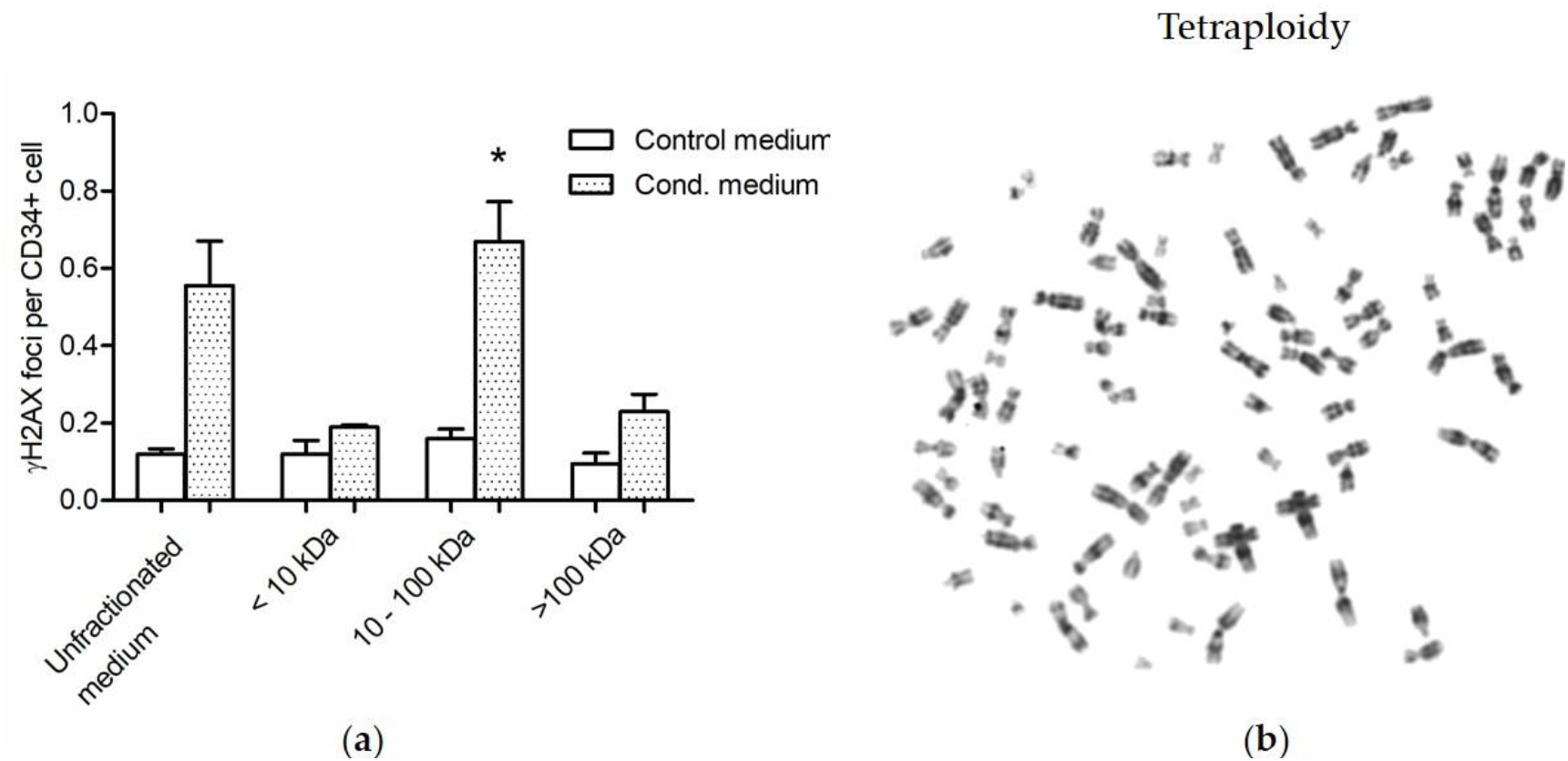
Radiation-induced bystander effects in CD34+ cells grown in distinct molecular size fractions of medium conditioned by 2 Gy irradiated mesenchymal stromal cells (MSC) and un-/ fractionated control medium. (**a**) γH2AX foci levels in CD34+ cells grown in (I) < 10 kDa, (II) 10 – 100 kDa and (III) > 100 kDa fractions of MSC conditioned medium and in un-/fractionated control medium. * *p* = 0.0068 [Kruskal-Wallis test] and *p* = 0.0286 [Wilcoxon two-sample test] when compared to numbers of γH2AX foci in CD34+ cells grown in (I) < 10 kDa and (III) > 100 kDa fractions or in un-/fractionated control medium. (**b**) Exemplary tetraploidy of a CD34+ cell grown in the (II) 10 – 100 kDa fraction of MSC conditioned medium.

**Table 1.**
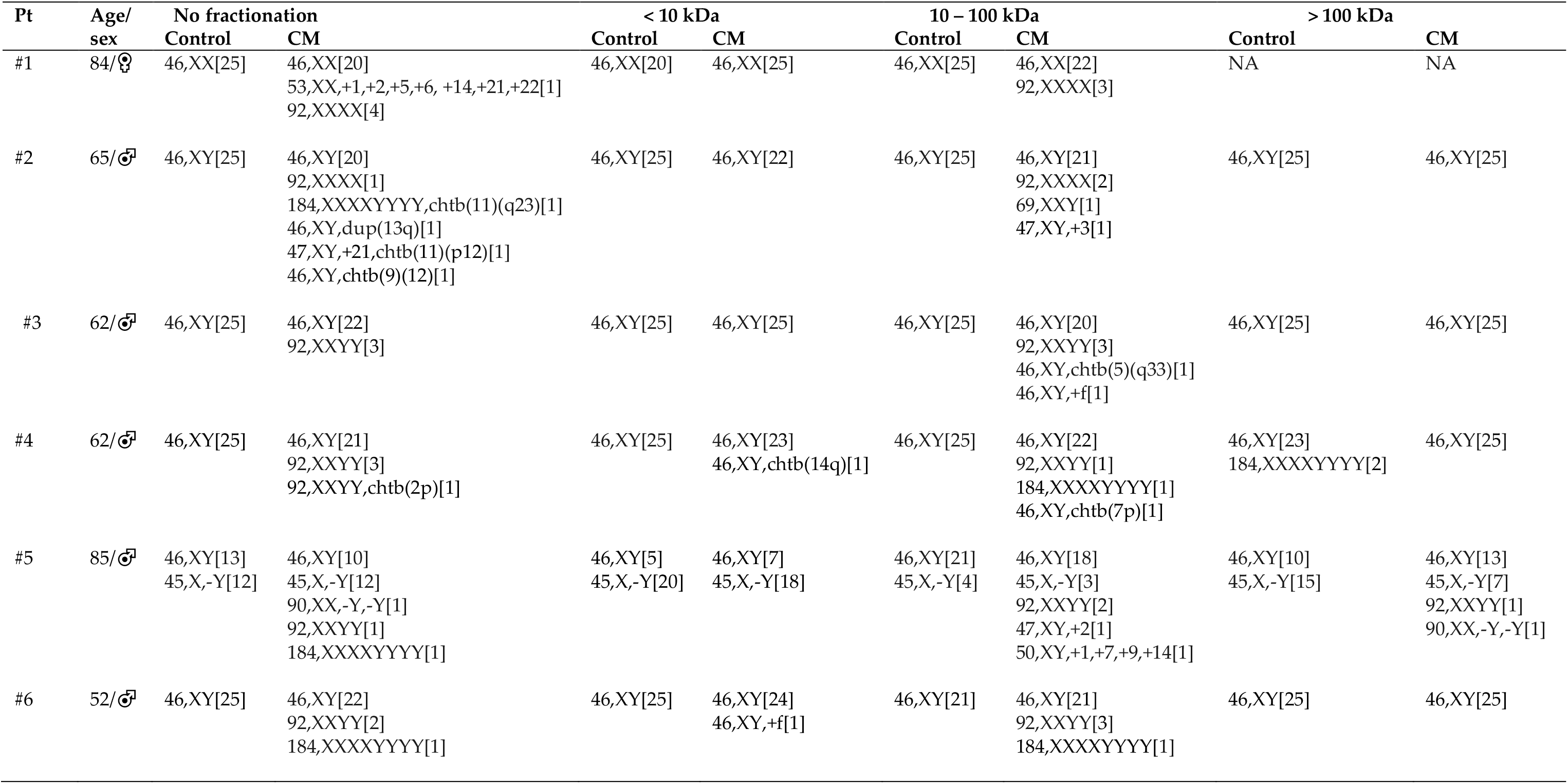
Radiation-induced bystander effects in CD34+ cells grown in un-/fractionated medium conditioned by 2 Gy irradiated mesenchymal stromal cells. CM, conditioned medium; NA, not assessed.

### 3.2. Chromosomal instability in human CD34+ cells

Metaphases were analyzed in human CD34+ cell samples (6 patients; ∑ 46 samples) expanded for 3 days in native medium followed by culture for 3 days in un-/fractionated MSC conditioned or un-/fractionated control medium, respectively (Figure 1b, Table 1). Increased numbers of aberrant metaphases (general *p* = 0.0007 [Kruskal-Wallis test]; pairwise comparison each *p* = 0.0022 [Wilcoxon two-sample test]) were detected in CD34+ cells grown in the (I) 10 – 100 kDa fraction of MSC conditioned medium when compared to numbers of aberrant metaphases in CD34+ cells grown in (II) < 10 kDa and (III) > 100 kDa fractions of MSC conditioned medium or in un-/fractionated control medium. More precisely, distinct chromatid breaks (chtb), e.g., chtb(5q) and chtb(7q) as well as aneuploidies, e.g., tetraploidies and octaploidies, were observed in CD34+ cells grown in the (II) 10 – 100 kDa fraction of MSC conditioned medium. In addition, distinct chtb, e.g., chtb(2), chtb(9) and chtb(11) as well as aneuploidies, e.g., tetraploidies and octaploidies, were observed in CD34+ cells grown in unfractionated MSC conditioned medium. It has to be noted, that loss of chromosome Y in #5 is a common finding in elderly men occurring at a frequency of 5 – 10% [25, 26]. Further, few chromosomal aberrations, e.g., chtb(14q) and aneuploidies, e.g., tetraploidies, were detected at very low frequencies in (I) < 10 kDa and (III) > 100 kDa fractions of MSC conditioned medium, which might be due to limitations in accuracy of the filtration process.

Finally, heat inactivation of unfractionated MSC conditioned medium and unfractionated control medium (2 patients, ∑ 4 samples) resulted in reduced proliferation of CD34+ cells. Here, all evaluable metaphases displayed a normal karyotype.

## 4. Discussion

Genotoxic bystander signals released from irradiated human MSC may induce DNA damage and CIN in human HSPC potentially initiating MN. While increased DNA damage and CIN are readily inducible in human CD34+ cells by exposure to MSC conditioned medium, the genotoxic bystander signals in MSC conditioned medium remain largely uncharacterized yet. Therefore, our study was designed to investigate the molecular features of bystander signals in terms of molecular weight and potential protein characteristics. For this purpose, approximate (I) < 10 kDa, (II) 10 – 100 kDa and (III) > 100 kDa fractions of MSC conditioned medium were generated for co-culture experiments in healthy human CD34+ cells of the same donors.

Increased numbers of γH2AX foci were detected in CD34+ cells grown in the (II) 10 – 100 kDa fraction of MSC conditioned medium when compared to low numbers of γH2AX foci in CD34+ cells grown in (I) < 10 kDa and (III) > 100 kDa fractions of MSC conditioned medium or in un-/fractionated control medium. As γH2AX foci are a marker of DSB, our data are in line with similarly increased numbers of chtb detected in CD34+ cells grown in the (II) 10 – 100 kDa fraction of MSC conditioned medium. Importantly, chtb may activate oncogenes or inactivate tumor suppressor genes, respectively, thus providing a potential mechanistic link to the initiation of MN [27].

Further, increased numbers of aberrant metaphases were observed in CD34+ cells grown in the (II) 10 – 100 kDa fraction of MSC conditioned medium when compared to low numbers of aberrant metaphases in CD34+ cells grown in (I) < 10 kDa and (III) > 100 kDa fractions of MSC conditioned medium or in un-/fractionated control medium. In particular, the number of tetraploidies was increased in the (II) 10 – 100 kDa fraction of MSC conditioned medium. Generally, tetraploidies may occur by chromosomal non-disjunction during mitosis or cytokinesis failure [28]. Further, tetrapolidies are found in about 1% of AML but 13% of t-AML cases [29]. Hence, our finding of increased tetraploidies in CD34+ cells grown in the (II) 10 – 100 kDa fraction of MSC conditioned medium suggests a mechanistic link to the initiation of MN. Although tetraploidies occurred at very low frequency in CD34+ cells grown in control medium, this result is not contradictory to our interpretations but indicates that tetraploidies may randomly occur *in vitro* during the proliferation process itself.

Finally, heat inactivation of unfractionated MSC conditioned and control medium resulted in reduced proliferation of CD34+ cells, which all demonstrated regular karyotypes. Thus, RIBE mediators have a temperature-sensitive structure, supporting the notion that the three-dimensional conformation of macromolecules, such as the native tertiary structure in proteins, confers specifically to the genotoxic effects in the (II) 10 – 100 kDa fraction of MSC conditioned medium instead of the sheer presence of mediating macromolecules.

Our study may raise the question for the impact of ROS and NO as potential RIBE mediators in the 10 – 100 kDa fraction of MSC conditioned medium. Considering that ROS and NO are rather short-lived mediator molecules, there might be no major impact of MSC released ROS and NO on detected RIBE in CD34+ cells in our experiments. More likely, hitherto unknown mediators with a longer half-life may increase ROS and NO in exposed CD34+ cells grown in MSC conditioned medium [20].

## 5. Conclusions

In conclusion, our data demonstrate that substantial genotoxic bystander signals mainly localize in the (II) 10 – 100 kDa fraction of MSC conditioned medium and that these signals are heat-sensitive. Based on these biochemical properties, we postulate proteins as RIBE mediators, which should be further analyzed by an in-depth proteome analysis of the corresponding fraction. Ultimately, it has the potential to uncover the identity of key bystander signals, which is fundamental for the development of next-generation anti-leukemic drugs.

## Acknowledgments

This research was funded by Deutsche José Carreras Leukämie-Stiftung, DJCLS 14 R/2017.

## Author Contributions

Conceptualization, H.D.P. and O.D.; methodology, H.D.P and O.D.; software, V.K. and A.F.; validation, V.K., A.F. and H.D.P.; formal analysis, C.W.; investigation, V.K., S.B. and A.F.; resources, H.K., A.D., A.J., M.B., W.S., A.F. and W.-K.H.; data curation, V.K., A.F. and H.D.P.; writing—original draft preparation, H.D.P; writing—review and editing, H.D.P., O.D., J.F., W.S., A.F. and W.-K.H.; visualization, H.D.P.; supervision, A.F. and W.-K.H.; project administration, H.D.P.; funding acquisition, H.D.P. and A.F. All authors have read and agreed to the published version of the manuscript.

## Abbreviations

The following abbreviations are used in this manuscript:

chtb: chromatid breaks
CIN: chromosomal instability
DSB: DNA double-strand breaks
HSPC: human hematopoietic stem and progenitor cells
MN: myeloid neoplasms
MSC: mesenchymal stromal cells
NO: nitric oxide
PBS: phosphate-buffered saline
RIBE: radiation-induced bystander effects
ROS: reactive oxygen species
t-MN: therapy-related MN

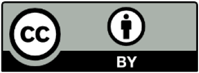 © 2020 by the authors; licensee MDPI, Basel, Switzerland. This article is an open access article distributed under the terms and conditions of the Creative Commons by Attribution (CC-BY) license (http://creativecommons.org/licenses/by/4.0/).

